# LSTrAP-Kingdom: an automated pipeline to generate annotated gene expression atlases for kingdoms of life

**DOI:** 10.1101/2021.01.23.427930

**Authors:** William Goh, Marek Mutwil

## Abstract

**Summary:** There are now more than two million RNA sequencing experiments for plants, animals, bacteria and fungi publicly available, allowing us to study gene expression within and across species and kingdoms. However, the tools allowing the download, quality control and annotation of this data for more than one species at a time are currently missing. To remedy this, we present the Large-Scale Transcriptomic Analysis Pipeline in Kingdom of Life (LSTrAP-Kingdom) pipeline, which we used to process 134,521 RNA-seq samples, achieving ~12,000 processed samples per day. Our pipeline generated quality-controlled, annotated gene expression matrices that rival the manually curated gene expression data in identifying functionally-related genes.

**Availability and implementation:** LSTrAP-Kingdom is available from: https://github.com/wirriamm/plants-pipeline and is fully implemented in Python and Bash.

## 1. Introduction

When first developed, the main application of RNA sequencing (RNA-seq) was for performing differential gene expression between samples (Stark *et al.*, 2019). The decreasing cost of sequencing and computational power has driven the explosive growth of RNA-seq data (https://www.genome.gov/about-genomics/fact-sheets/DNA-Sequencing-Costs-Data), evident in the increase in the amount of open access data on Sequence Read Archive (SRA) from 100 Terabases in 2011 (Kodama *et al.*, 2012), to more than 10,000 Terabases in 2020 (https://www.ncbi.nlm.nih.gov/sra/docs/sragrowth/). The RNA-seq data is especially abundant for animals (1,774,720 samples), followed by plants (173,663), fungi (53,712), and bacteria (50,615), at the time of writing of this manuscript. The RNA-seq data allows coexpression analyses useful in gene function prediction and to supplement integrated multi-omics analyses (Usadel *et al.*, 2009; Rhee and Mutwil, 2014). Furthermore, when combined with genomic information, comparative transcriptomic analyses across species allow the study of the function and evolution of genes from the perspective of gene expression (Ferrari *et al.*, 2019; Wen Tan and Mutwil, 2019; Ferrari *et al.*, 2020; Ng *et al.*, 2019; Ferrari and Mutwil, 2019; Lim *et al.*, 2020).

Thus, the available gene expression data provides an untapped opportunity to study gene function in the kingdom of life, but user-friendly tools to download, quality control, annotate and collate the RNA-seq data for more than one species at a time are currently not available. To remedy this paucity, we constructed an open-source pipeline, Large-Scale Transcriptomic Analysis Pipeline in Kingdom of Life (LSTrAP-Kingdom), that allows rapid generation of kingdom-wide expression atlases. We used our automated pipeline to download 134,521 RNA-seq experiments for 116 species, achieving processing speed of ~12,000 samples per day, and show that the experiments can be annotated with a simple natural language processing pipeline that leverages organ ontology information. Finally, we show that the coexpression networks obtained by our pipeline perform as well as networks constructed from manually assembled matrices.

## 2 Automated retrieval of coding sequences, available experiments, and gene expression value estimation

To demonstrate our pipeline, we used *Viridiplantae* (taxid 33090), encompassing all green plants (Figure 1A). LSTrAP-Kingdom retrieves taxonomic IDs (taxid) for all species found under the clade of interest from NCBI Taxonomy API and SRA browser (Kodama *et al.*, 2012). Then, for each species, the pipeline retrieves a list of all available RNA-seq experiments found in the SRA database and links to coding sequence (CDS) files from Ensembl FTP directories (Kersey *et al.*, 2016). The pipeline then generates a tab-delimited species list table where the user can edit, add or remove the links to CDS files (Table S1). Only species with CDS links specified in the table will be further processed, allowing the user to define which species should be analyzed by the pipeline, and also to specify alternative sources of CDS files. For each species in the curated species list table, LSTrAP-Kingdom retrieves the CDS files, generates kallisto index file, downloads the available fastq files (with an option for a parallel download), and estimates the gene expression with kallisto (Bray *et al.*, 2016). LSTrAP-Kingdom will also download the SRA run tables for each species, containing the metadata of the RNA-seq experiments used to annotate the experiments (Table S2).

**Figure 1:**
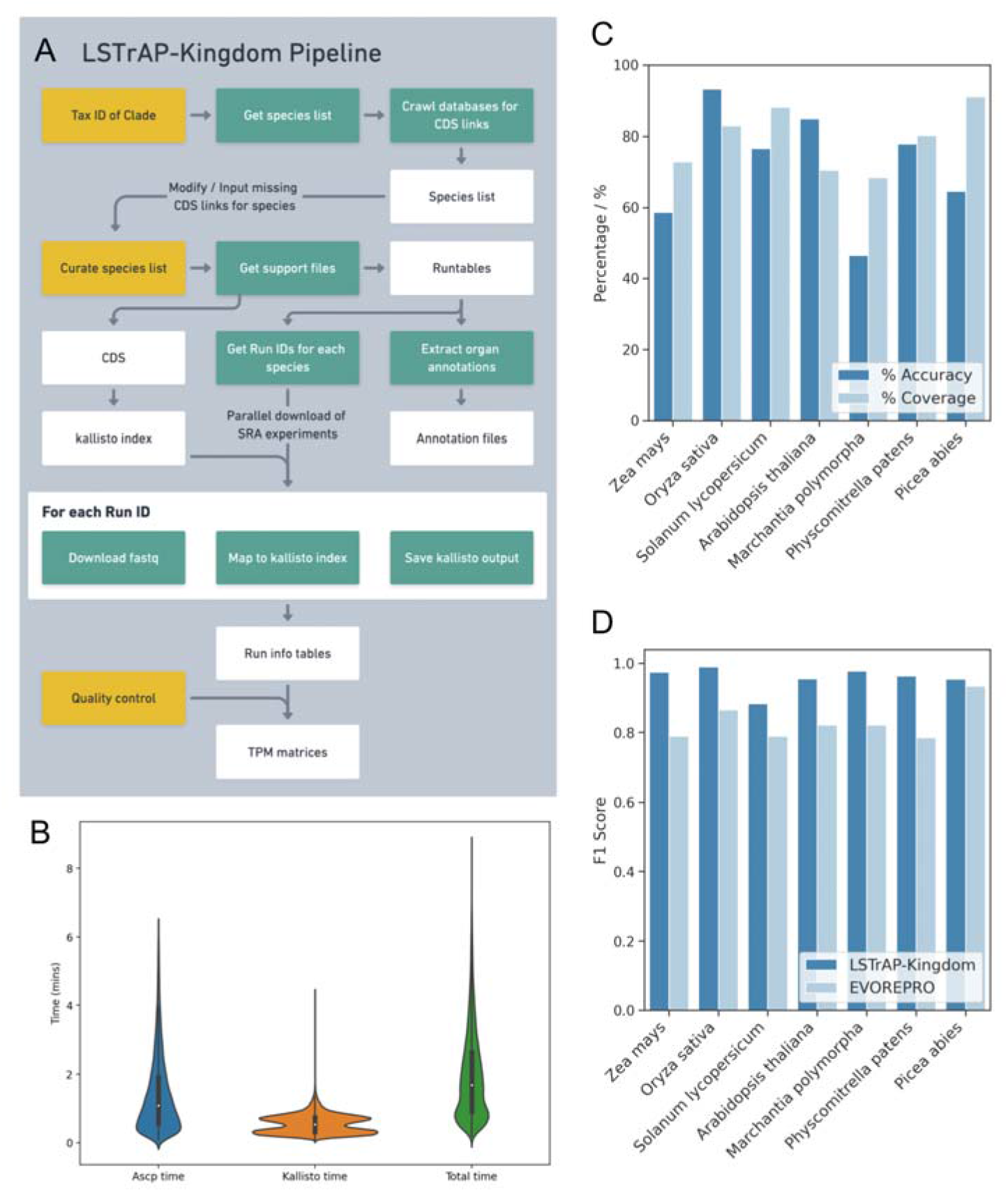
The LSTrAP-Kingdom pipeline. A) An outline of the pipeline. Yellow: user input; green: processes; white: output files. B) Distribution of time taken (in minutes) for ascp download, Kallisto processing, and total time for 35,020 samples from *Arabidopsis thaliana.* C) Percentage accuracy and coverage of LSTrAP-Kingdom annotation benchmarked against the manually annotated samples from the EVOREPRO study. D) F1 scores of coexpression networks generated by LSTrAP-Kingdom in predicting genes for biosynthesis of ribosomal proteins, benchmarked against the EVOREPRO dataset.

In our example, we selected 116 Archaeplastida species for download. With 24 parallel downloads, we could download ~12,000 samples per day, when the fastq file size was capped at 1GB (~20 million reads)(Table S3). For *Arabidopsis thaliana* (taxid 3702), the mean time taken for download, gene expression estimation with kallisto, and total processing is 1.41 minutes, 0.55 minutes, and 1.95 minutes, respectively (Figure 1B). To perform quality control of the downloaded RNA-seq, LSTrAP-Kingdom produces a table describing the RNA-seq data with two param eters: log10-normalized number of processed reads (LPR) and percentage of reads pseudo-aligned to the reference CDS (PPA)(Tan *et al.*, 2020). These parameters were used to visualize any outlier samples (Figure S1) and to recommend LPR and PPA thresholds to remove samples with a low number of reads or poor mapping statistics (Table S4). Samples accepted by the user (Figure S2) are used to compile a TPM matrix for each species.

## 3 Automated sample annotation

LSTrAP-Kingdom annotates the experiments by matching words found in the experiment metadata with any organ ontology in obo format. For the Archaeplastida kingdom, we used Plant Ontology’s (PO) anatomical entity domain to specify plant organs and tissues (Walls *et al.*, 2019). First, for each PO term (e.g., leaves, roots, seeds), the words in a PO term were stemmed (leav, root, seed), and the percentage of the stemmed words of each PO found in a given RNA-seq experiment description was calculated. Then, each RNA-seq sample was annotated with the PO term showing the highest percentage of matched words. If multiple equal matches were found, the PO term with most words was used to annotate the RNA-seq experiment.

We benchmarked the accuracy of our pipeline against manual annotation of RNA-seq data from 7 species in the EVOREPRO study (Table S5, S6)(Julca *et al.*, 2020). The proportion of samples that could be annotated range from 68% to 91% (Figure 1C). The accuracy ranged from 46.5% in *Marchantia polymorpha* to 93.3% in *Oryza sativa* (Figure 1C). The poor accuracy in *Marchantia polymorpha* was due to the description of the RNA-seq samples in the run table not using standard nomenclature (Table S5). For example, some samples were annotated as “sporelings”, a developmental stage of bryophytes not represented in PO. Thus, a more comprehensive PO database and a standardized sample description nomenclature will improve future automated sample annotation. Thus, our automated pipeline can be used to rapidly annotate thousands of RNA-seq samples (Table S7).

## 4 Evaluation of coexpression networks generated

To measure the performance of the coexpression networks generated by our pipeline, we evaluated how well our networks predict genes that are components of large and small ribosomal subunits (Supplementary Figure S3), as was done in a previous study (Hew *et al.*, 2020). Briefly, each ribosomal protein was functionally annotated by its coexpression neighborhood functions, at a given Pearson correlation coefficient (PCC) threshold and percentage of coexpression neighborhood required to annotate the function. For example, a gene is annotated as a ribosomal protein if at least 50% of it’s coexpression neighborhood at PCC>0.7 are ribosomal proteins. As a metric for the predictive power, we used F1-score, which is a harmonic mean between the precision and recall of the predictions of the coexpression network.

The F1-scores range from 0.8828 in *Picea abies* (taxid 3329) to 0.9896 in *Physcomitrium patens* (taxid 3218) for the coexpression networks generated by LSTrAP-Kingdom (Figure 1D), showing a strong predictive power of the generated coexpression networks. Furthermore, the coexpression networks generated in this study show slightly higher F1-scores than the networks obtained from the manually constructed matrices used in the EVOREPRO study (Julca *et al.*, 2020). This demonstrates the usefulness of LSTrAP-Kingdom for kingdom-wide expression analysis.

## Data Availability Statements

The data underlying this article are available in the article and in its online supplementary material. The RNA-sequencing data is publicly available from NCBI SRA.

## Supplementary materials

**Table S1. Species list for plants identified by Viridiplantae taxonomy ID (33090).** The table shows the taxonomic ID, species name, RNA sample count (total), RNA sample count (Illumina), and a link to a CDS file found for each species. Links which are do not belong to ‘ensemblegenomes’ were manually added.

**Table S2. SRA runtables used for annotating the seven species.**

**Table S3. Logfile of download times for each species in the batch.** The table shows the species number, taxonomic ID, the attempt number (failed downloads are retried two times), total number of downloaded fastq files, total number of failed downloads, number of downloaded files in a current attempt and time taken for the current attempt.

**Table S4. Quality control thresholds (A) recommended by LSTrAP-Kingdom and (B) edited by the user.** The pipeline automatically recommends cutoff thresholds for log_10_ processed and % pseudoaligned and shows how many fastq files pass the given threshold. The user can adjust the cutoffs to either increase or decrease the stringency.

**Table S5. Comparison of LSTrAP-Kingdom’s annotation against EVOREPRO’s annotation.** The table shows the taxonomy ID, species name, the fastq file name (run ID), EVOREPRO annotation of the sample (comprising ten categories: root, seeds, stem, leaf, flower, male, female, root meristem, apical meristem, spore), PO term predicted by the pipeline and outcome of the prediction (true or false). Since the EVOREPRO annotation is more general (e.g., female) than the PO terms (e.g., plant egg cell), we indicated where the seemingly incorrect predictions are correct in the ‘modified annotation’ column.

**Table S6. Mapping of EVOREPRO annotation terms to the related PO terms.** The table used to translate the more specific PO terms to the more general EVOREPRO sample annotations, as done in the ‘modified annotation’ column in Table S5.

**Table S7. LSTrAP-Kingdom sample annotation for seven species**. The species are *Marchantia polymorpha, Physcomitrium patens, Picea abies, Arabidopsis thaliana, Solanum lycopersicum, Oryza sativa* and *Zea mays.*

**Figure S1.**
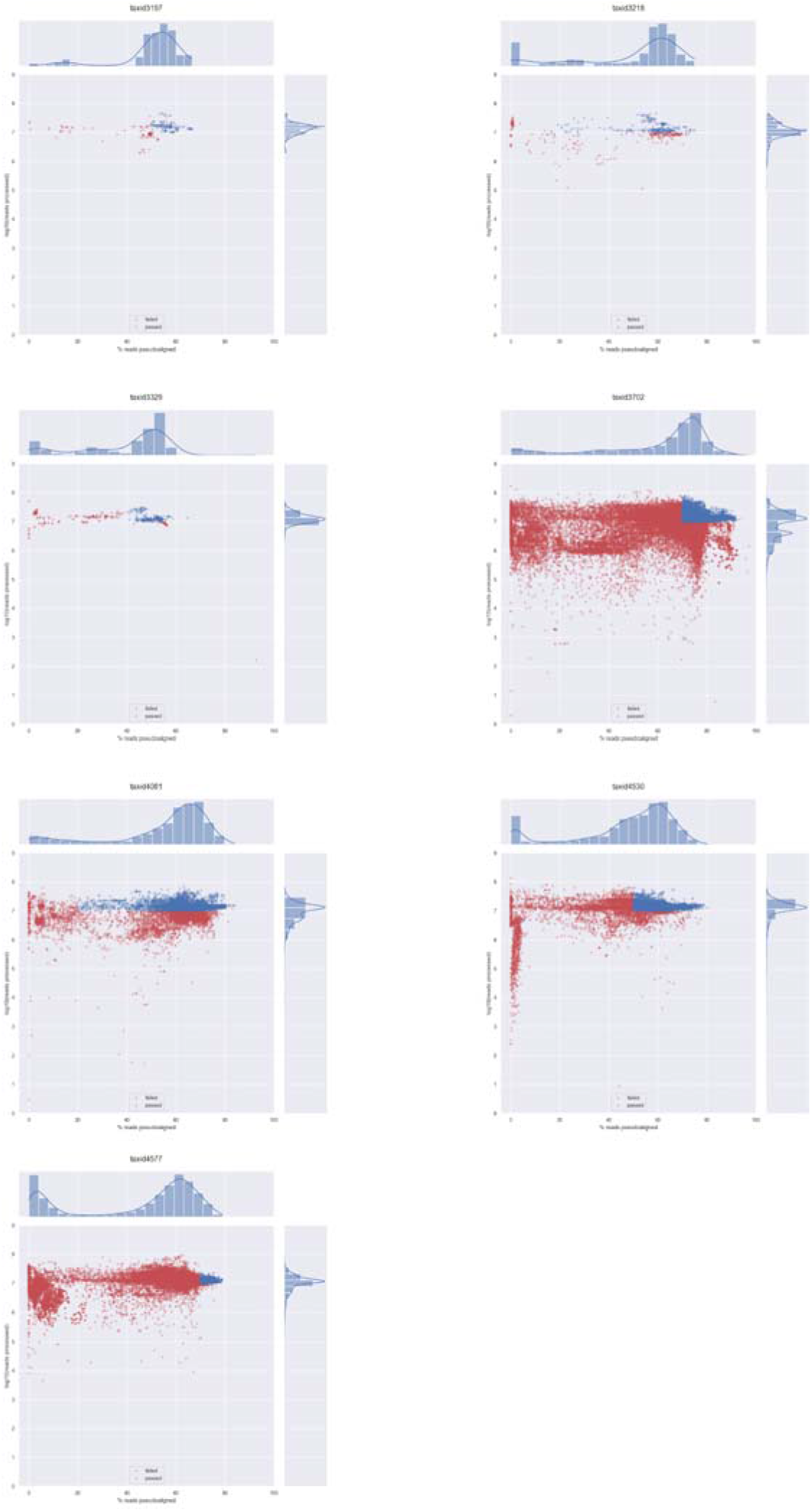
Quality control visualization with the LSTrAP-Kingdom automatically suggested thresholds. Each point represents a fastq experiment, where a blue or red point represents experiments that passed or failed the thresholds, respectively. The x- and y-axis represent the % of pseudoaligned reads and log_10_ processed reads, respectively.

**Figure S2.**
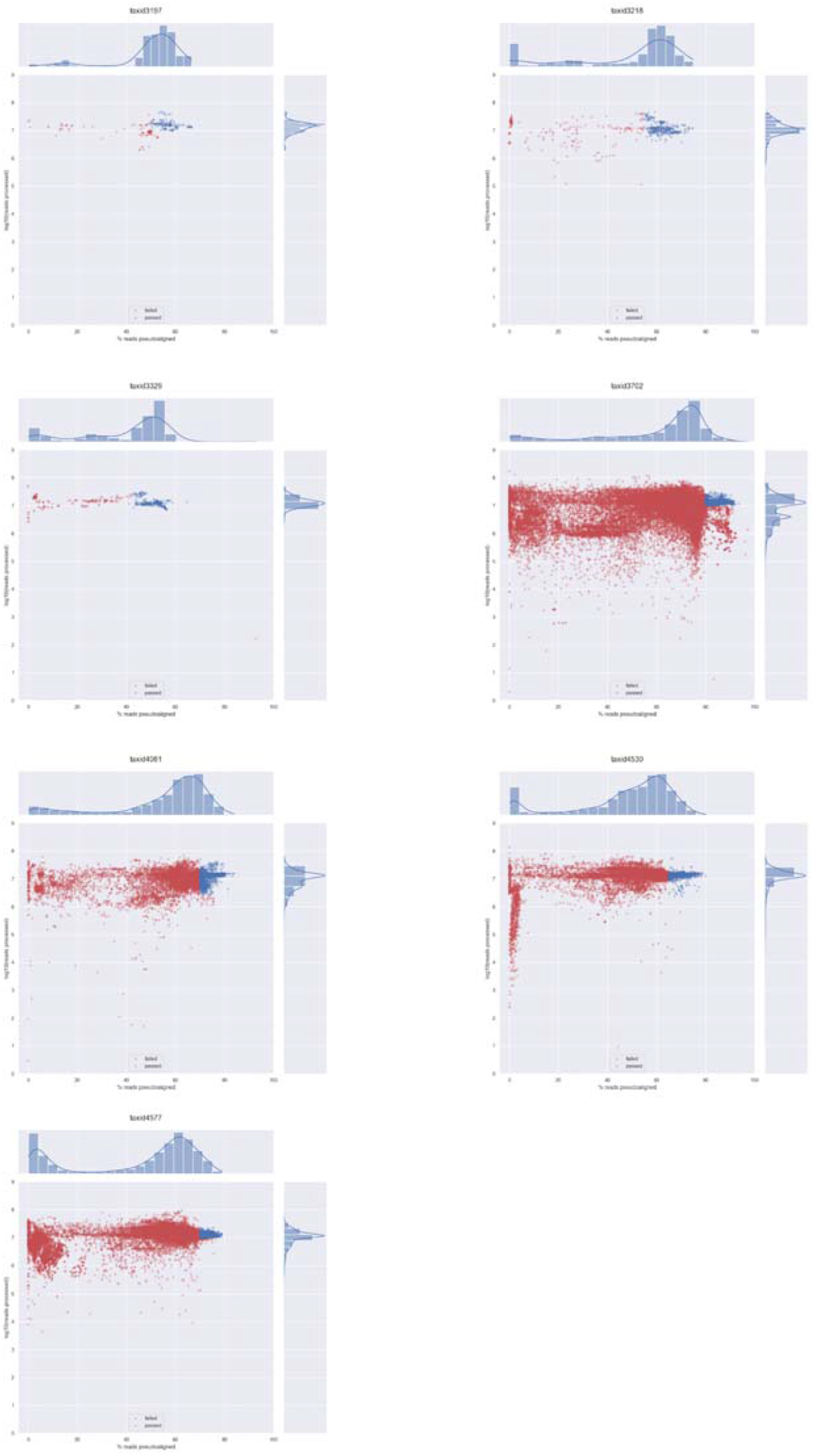
QC visualization with user-defined thresholds. Thresholds from Table S4B are used in this figure.

**Figure S3:**
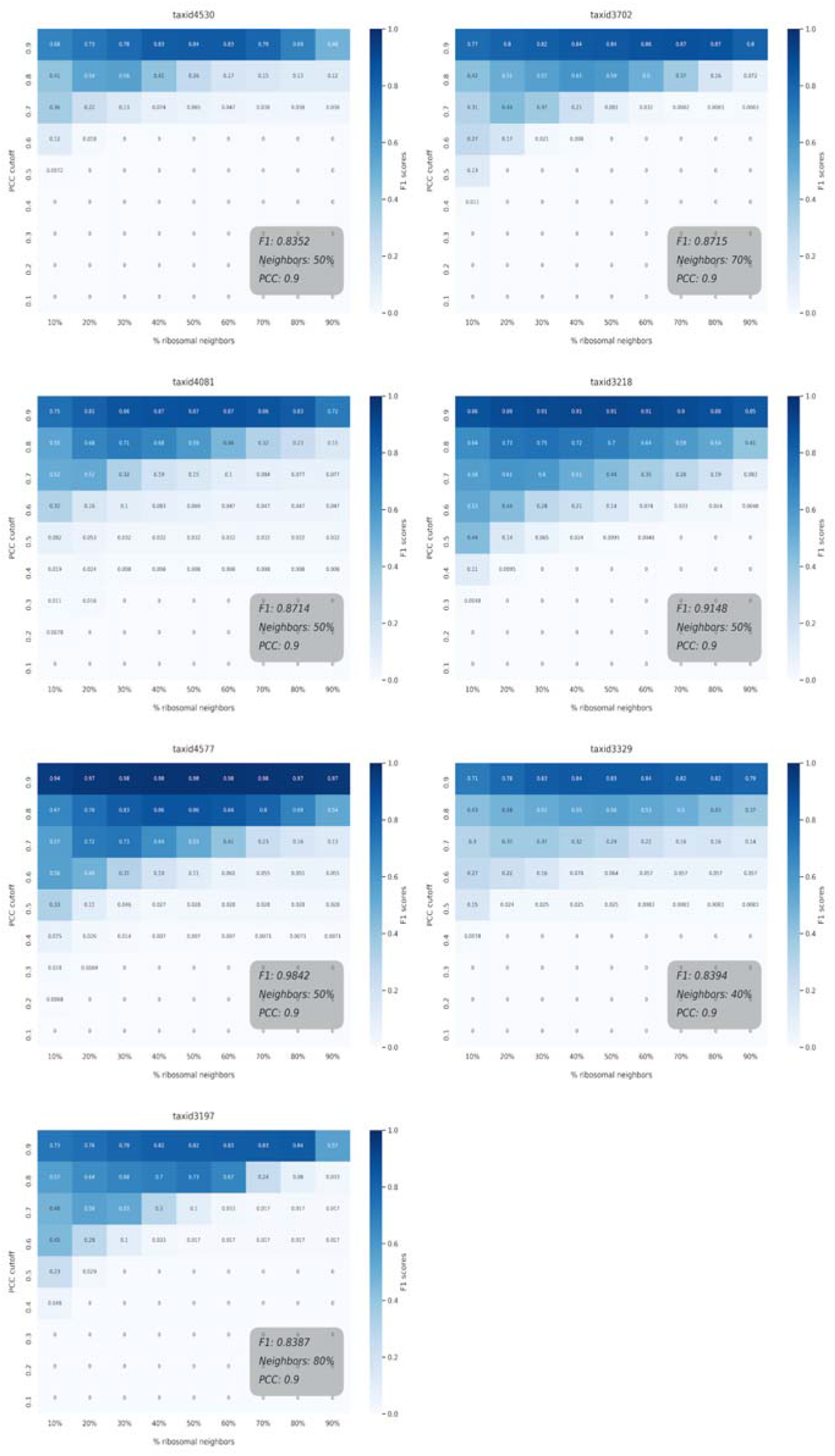
Heatmaps of F1 scores for LSTrAP-Kingdom. The heat maps indicate the F1-score for each used pair of PCC and the percentage of ribosomal neighbors thresholds.

